# Genome-wide temporal gene expression reveals a post-reproductive shift in the nematode *C. briggsae*

**DOI:** 10.1101/2024.06.04.597319

**Authors:** Wouter van den Berg, Bhagwati P Gupta

## Abstract

*C. briggsae* offers a robust system for comparative investigations of genetic pathways that affect physiological processes. One key process, reproduction, significantly impacts longevity due to its high energetic cost, which limits resources for somatic maintenance. Long-lived mutants often exhibit reproductive impairments, and studies in *C. elegans* have demonstrated that germline mutations and complete germline removal can promote longevity, underscoring the link between reproduction and aging. We are interested in identifying genes and biological processes affected during the reproductive and post-reproductive periods in *C. briggsae*. To achieve this, we conducted whole-genome transcriptome profiling on animals at various adult stages. analysis of differentially expressed (DE) genes revealed that the majority were downregulated during the reproductive period. Interestingly, this trend reversed post-reproduction, with three-quarters of the genes upregulated—a phenomenon we termed the ‘reproductive shift’. A similar analysis in *C. elegans* also showed a downregulation bias during the reproductive period, but the reproductive shift was absent. Further examination of *C. briggsae* DE genes showed enrichment in processes related to the matrisome, muscle development and function during the reproductive period. Post-reproductive downregulated genes were enriched in DNA damage repair, stress response, and immune response. Additionally, terms related to fatty acid metabolism, catabolism, and transcriptional regulation exhibited complex patterns, with different biological processes being up or downregulated between the reproductive and post-reproductive stages. Overall, our transcriptomic data provides a valuable resource for cross-sectional comparative studies of reproductive and post-reproductive changes in nematodes. Additionally, the findings prompt similar studies in other animal models thereby advancing our understanding of genetic pathways affecting reproduction and aging.

## INTRODUCTION

Life reproduces life, and the process of achieving this is energetically costly. Particularly for females, reproducing is known to have a negative relationship with the lifespan of an animal (Duggal, 1978; Maklakov and Immler, 2016; Partridge et al., 2005). As the soma deteriorates over time, few processes can be said not to be affected by aging. The process of reproduction is intimately involved with and affected by aging: germ cell quality reduces in older organisms, leading to birth defects and decreasing reproductive success (Brieno-Enriquez et al., 2022; Chu et al., 2023). Other processes, such as menopause in several mammal species, put a stop to reproduction at an older age, instigating a post-reproductive life stage that is the topic of much scientific speculation (Croft et al., 2015; Gems and Kern, 2022; Monaghan and Ivimey-Cook, 2023; Takahashi et al., 2016). Hermaphroditic nematodes also have an extensive post-reproductive lifespan, although likely for different reasons (Scharf et al., 2021). In these animals, self-fertile reproduction is limited by the self-sperm supply, although by mating with males the reproductive period can be extended until aging-related impairments become more relevant (Ward and Carrel, 1979).

As a member of the *Caenorhabditis* genus, *C. briggsae* has been studied as an independent model as well as a satellite model to the widely studied *C. elegans* (Alliance of Genome Resources, 2024; Gupta et al., 2007). Investigating the physiological processes and expression of orthologs and homologs in these two species allows for the correct interpretation of genetic and functional conservation of genes. Although *C. briggsae* is morphologycially similar to *C. elegans*, there are notable differences (Gupta et al., 2007; Jhaveri et al., 2023). While both nematode species are androdioecious, the hermaphroditic reproductive mode of *C. briggsae* evolved separately from that of *C. elegans* (Guo et al., 2009; Kiontke et al., 2004). *C. briggsae*-*C. elegans* comparative studies allow for understanding traits that are conserved versus species-specific as well as the molecular basis of these traits.

As a leading invertebrate model, *C. elegans* has facilitated comparisons across organisms (Gems and Partridge, 2013; Girardot et al., 2006; McCarroll et al., 2004; van der Goot et al., 2012; Wang et al., 2014). Several longitudinal studies have examined transcriptome patterns during post-developmental time in *C. elegans* (Budovskaya et al., 2008; Golden et al., 2008; Tarkhov et al., 2019). In both *C. elegans* and *C. briggsae*, oocytes are fertilized by self-sperm, which is the limiting factor on reproductive capacity. Unmated hermaphrodites maintain oogenesis and yolk protein production post-reproduction, but these are impotently expelled (Hansen and Schedl, 2006; Kern et al., 2023). Mating allows worms to extend their brood size and reproductive span which normally ends sometime between day 6 and day 9 (Scharf et al., 2021).

Many longitudinal studies in *C. elegans* have focused on transcriptome changes in the context of aging or looked at the long-term effects of experimental interventions such as exposure to chemicals or in knockout models (e.g., see (Li et al., 2019; Schmeisser et al., 2013)). These studies have revealed gene expression changes at different adult stages including the reproductive span, but none have specifically focused on discussing reproductive to post-reproductive expression changes. Reinke et al. identified germline genes in *C. elegans* through differential transcriptomics in germline-impaired mutants (Reinke et al., 2004; Reinke et al., 2000). These sets include genes directly involved in reproduction, as well as downstream factors and transcription factors such as *lin-35*, E2F heterodimer factors *efl-1*, and *dpl-1*, and *spe-44*, which regulate reproduction related processes (Ceol and Horvitz, 2001; Chi and Reinke, 2006; Kulkarni et al., 2012; Mikeworth et al., 2023; Ragle et al., 2022). Evolutionary conservation of germline genes within other nematodes has been studied (Artieri et al., 2008; Cutter and Ward, 2005), finding rapid evolution of coding and regulatory sequences compared to somatic genes. However, genome-wide expression studies are needed to facilitate a better understanding of differences and similarities of germline processes and reproduction in general.

Besides the role of aging in the reproductive capacity of nematodes, longitudinal studies have determined that many different physiological processes change and become disrupted with age including lipid and protein metabolism, mitochondrial function and collagen metabolism (Golden et al., 2008). Aging affects the expression of extracellular matrix proteins, collectively referred to as the matrisome, which are constantly adjusted to maintain homeostasis but decline after reproduction(Ewald, 2020). Longitudinal studies that focus on aging often include timepoints of the adult lifespan that cover the reproductive as well as the post-reproductive period, but the effects of this transition have been ignored despite extensive interactions of many processes with reproduction. Lipid metabolism functions in a seesaw manner, balancing resources between survival and reproductive investment (Hansen et al., 2013). Movement and muscle function assays in *C. elegans* have also reported a gradual decline as age progresses (Herndon et al., 2002). Moreover, movement velocity decreases with age, and the decline becomes steeper between day 5 and day 10 as the reproductive period ends (Hahm et al., 2015; Koopman et al., 2020).

In this study, we sought to determine genome-wide expression patterns of *C. briggsae* in a longitudinal manner, spanning the reproductive to the post-reproductive period. Our analysis showed a ‘reproductive shift’ in the transcriptome, leading us to investigate affected processes, involvement of germline genes, and reproduction-related transcription factors. Our findings included post-reproductive upregulation of the matrisome and muscle processes, and downregulation of DNA repair, immunity and stress response. Results were compared with transcriptome data of *C. elegans*, which revealed the reproductive shift transcriptome pattern to be stronger in *C. briggsae*. The post-reproductive expression changes of matrisome and several germline genes appear to be partially conserved. We also showed that mating affects gene expression. Finally, a comparison of downstream genes of a set of germline-related transcription factors revealed both conserved and species-specific expression trends. Overall, our findings serve as a valuable resource to facilitate the study of biological processes affecting reproductive and post-reproductive states of *C. briggsae* and enable comparative studies with other nematodes. Our work provides a foundation for future research exploring conserved mechanisms and their implications across different animal models.

## METHODS

### Worm culture maintenance

*C. briggsae* AF16 culture was obtained from the *Caenorhabditis* Genetic Center (CGC). *Cbr-glp-4(v473)* was kindly provided by Ronald Ellis (Rowan University). Worms were grown on Nematode Growth Medium (NGM) plates seeded with *E. coli* OP50 bacteria and maintained at 20°C, unless otherwise stated. For reproductive span determination, adult hermaphrodites were allowed to lay eggs for one hour to generate a synchronized cohort of parental hermaphrodites. Parental hermaphrodites were transferred, as young non-gravid adults, to individual plates and then transferred to new plates every 24 hr. Transfers continued until the cessation of egg-laying. Progeny on individual plates were counted two days after parental hermaphrodites were transferred.

### RNA isolation and sequencing

For RNA isolation, age-synchronized populations were obtained by two successive rounds of bleaching of young gravid adults by standard methods using NaOCl and NaOH (Stiernagle, 2006). Animals were left to pass through larval stages and live to specified ages: day 1, 3, 6, 9 of adulthood. Every second day, adults and larvae were column separated by washing in an upright 15 mL tube filled with M9 for 5 minutes to achieve size separation, after which the water column was pipetted off to remove larvae. This was repeated at least 3 more times to increase removal of larvae. Worms were pipetted on fresh seeded plates, and remaining larvae were removed with a platinum worm pick and burned. Worm populations at the specified age were washed in the same way, frozen at −80°C followed by RNA isolation.

RNA was isolated using Trizol-reagent (Sigma, USA, Catalog Number T9424), chloroform, and isopropanol (Chomczynski & Sacchi 1987). The quality of total purified RNA was confirmed using Nanodrop 1000 bioanalyzer (Thermofisher). cDNA libraries were constructed from 100–200ng RNA using an Illumina-specific commercial kit (TruSeq RNA Sample Preparation Kit v2, Set A, Catalog number RS-122-2001). RNA sequencing was carried out using an Illumina NovaSeq PE100 system at Génome Québec. For the day 1, 3 and 6 age categories, three biological replicates were used, consisting of full large plates of worms. For the day 9 age category, two biological replicates were used due to one sample not passing quality control. Paired-end reads were obtained for each cDNA library. Sequencing adapters were used with the following sequences: read 1 “AGATCGGAAGAGCACACGTCTGAACTCCAGTCAC”; read 2” AGATCGGAAGAGCGTCGTGTAGGGAAAGAGTGT”.

Sequence data were extracted in FASTQ format and used for mapping as described next. RNA-sequencing was performed with the following software and settings: MultiQC was used to aggregate multiple results into a single report (Ewels et al., 2016). Reads were aligned to the genome using RNA STAR, set to paired-end sequences (Dobin et al., 2013). The reference genome used was *C. briggsae* CB4 (Ross et al., 2011), in combination with WS279 annotation obtained from Wormbase. To count hits per gene, *featureCounts* was used with default settings for paired-end reads (Liao et al., 2014). Average input read counts were 33.0 M per sample (range 22.7 M to 39.2 M) and average percentage of uniquely aligned reads was 87.1% (range 85.8% to 88.1%). Differential gene expression analysis was performed with DESEQ2 in a pairwise manner (Love et al., 2014). Data based on individual biological replicates was merged into single age-based sets. In each comparison, the earlier timepoint was used as baseline.

The day 1 samples of the first and second sequencing runs were compared and found to be negligibly different. Further analyses involving the day 9 samples of the second sequencing run were done in comparison with the day 1, 3 and 6 samples of the first sequencing run. Genes were considered significantly differentially expressed on the later timepoint of a pairwise comparison for adjusted p-values <0.01. For those analyses where log2fold cutoff values were applied, limits were set for log2fold changes of −/+ 1. Figure 1 graphs were produced by DESEQ2 (Love et al., 2014).

**FIGURE 1.**
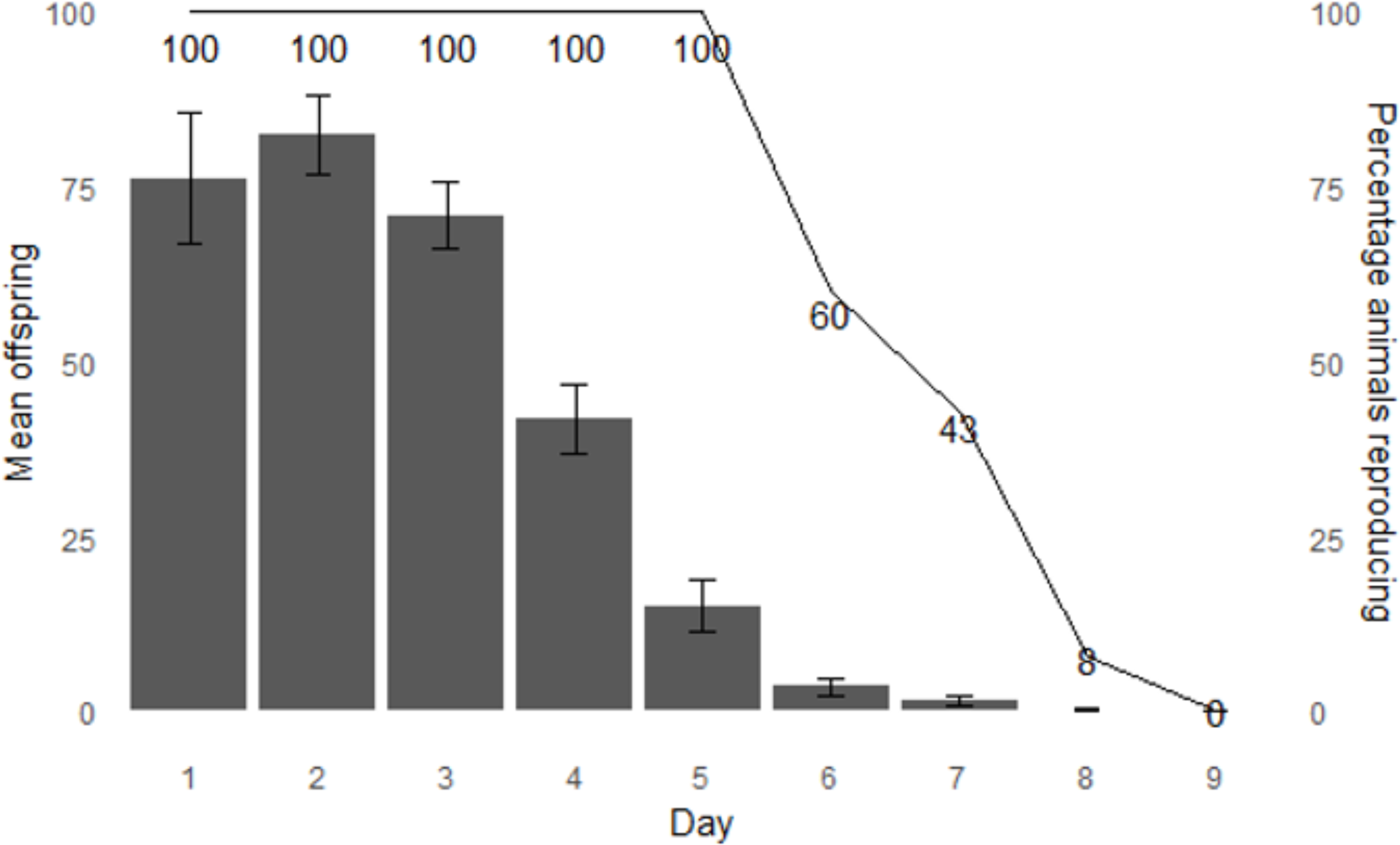
Reproductive span and brood size of *C. briggsae* AF16 unmated hermaphrodites. Bars display mean number of viable eggs laid by unmated *C. briggsae* hermaphrodites per day of adulthood. Error bars display standard error of the mean. Line displays the percentage of animals that remain reproductively active. N = 23.

### Gene annotation

Gene ontology (GO) and Kyoto Encyclopedia of Genes and Genomes (KEGG) analysis was performed on all expression subsets within the dataset of *C. briggsae* genes. *C. elegans* homologs of *C. briggsae* genes were determined using custom scripts in Perl, based on orthology data in the databases Inparanoid 8 (https://inparanoidb.sbc.su.se/), Wormbase-Compara, OrthoMCL (https://orthomcl.org/orthomcl/app), OMA (https://omabrowser.org/oma/home/) and Hillier et al. (Hillier et al., 2007). Analysis was done on both the *C. briggsae* genes and *C. elegans* orthologs of the equivalent sets of orthologous genes, i.e. genes without known homologs between the two species were excluded from annotation analysis. GO analysis was performed through the 17.0 release of PANTHER at https://archive.pantherdb.usc.edu/ (Thomas et al., 2022). KEGG analysis was performed on *C. elegans* orthologs, using KEGG database (Kanehisa and Goto, 2000) version 106.0. The tool KOBAS-i was used to perform KEGG analysis in a high-throughput manner on the top 3000 genes with greatest expression change per dataset (Bu et al., 2021). P-value are provided in two formats: uncorrected and Benjamini & Hochberg FDR corrected. GO and KEGG enriched terms were grouped into categories, each represented by multiple annotation terms at most or all timepoints. This was further manually trimmed to seven major categories based on gene set overlap and physiological relatedness.

### C. elegans comparison

Aging-associated expression patterns in *C. briggsae* were compared to *C. elegans.* Published microarray and RNAseq datasets (Golden et al., 2008; Schmeisser et al., 2013) were used as comparative longitudinal expression data in *C. elegans*. Main expression comparisons were performed using the Schmeisser et al. dataset (Schmeisser et al., 2013) that describes RNA-seq analysis of *C. elegans* N2 hermaphrodites grown at 20°C, aged to day 1, 5, 10. These time points correspond to reproductive (day 1 and day 5) and post-reproductive (day 10) stages of animals and, therefore, compared with *C. briggsae* samples of similar reproductive (day 1 and day 6) and post-reproductive (day 9) stages. For the Golden et al. dataset, comparisons were done using expression values for day 1, day 5, and day 9 of adulthood (Golden et al., 2008).

### Experimental analysis of reproduction-related genes

*Cbr-glp-4(v473)* were bleach synchronized and grown at 25°C until L4, subsequently switched to 26°C to induce full sterility, and grown to day 2 and day 6 of adulthood. AF16 controls were grown alongside at the same temperatures but started 12 hours later due to developmental delay in *Cbr-glp-4(v473)*. For extended reproduction assays, AF16 hermaphrodites were mated to males for 48 hours from day 1 and day 7 of adulthood. Mating success was confirmed by observation of copulatory plug on all animals after ∼24 hours. After reaching the designated ages, populations were washed 1x in M9, 1x in water to get rid of bacteria. After washing worms were left in 18μl water, 50 μl RNAzol (MRC Inc) and 2 μl precipitation carrier (MRC Inc) were added. RNA was extracted using isopropanol and followed standard methods. Samples were generated in at least three biological replicates. Based on the transcriptome data, up-invert and down-invert genes were selected for representative measurements. cDNA synthesis was performed using a SensiFAST cDNA synthesis kit (BIOLINE) with equalized amounts of RNA between samples. SensiFAST SYBR Green (BIOLINE) quantitative RT-qPCR was performed using the BIORAD CFX-96 Real Time system and following the BIORAD-CFX Software manual. Gene expression levels were normalized to housekeeping gene *Cbr-iscu-1*. Primers used are listed in the Supplemental table 1.

### Data analysis and statistics

Wormbase (http://www.wormbase.org) was used as a resource for Information about worm genes and pathways (Sternberg et al., 2024). For RNA-seq experiments, DEG counts were tracked and graphed in Microsoft Excel. Line graphs of figure 4 were produced in R, using ggplot2 (Wickham, 2016). Venn Diagrams were produced using Venny (Oliveros, 2007-2015). A hypergeometric test using the Benjamini & Hochberg false discovery rate (FDR) correction was implemented at a significance level of 0.05. Significance of gene set overlap was determined by a one-sided Fisher’s exact test. RT-qPCR results were analyzed using CFX Maestro 3.1 software (Bio-Rad, Canada), from three technical replicates for each run using the comparative 2ΔΔCt method and significance assessed by one-way ANOVA or t-test.

## RESULTS

### Transcriptome profiling during reproductive and post-reproductive stages

We set out to study gene expression changes in *C. briggsae* hermaphrodites during reproductive and post-reproductive periods. To this end, whole-body transcriptomes were analysed by RNA sequencing, sampled from populations of unmated worms aged to day 1, 3, 6 and 9 of adulthood. Figure 1 shows the reproductive span of AF16 unmated worms. The majority of fertilized eggs were laid during the first three days. By day 6, egg laying was nearly complete as the fertilized egg output dropped significantly. By this time, most individuals were laying their last few eggs (<10), while some had already depleted their capacity. By day 9, unmated individuals had fully stopped reproducing. The average reproductive span was 5.9 +/− 0.31 days with mean life-time brood size of 353.8 +/− 47.7 (n = 9). We, therefore, consider day 1-3 as the early reproductive phase, day 3-6 as late reproductive phase, and 6-9 and beyond as post-reproductive. Thus, the first three time points of transcriptome analysis overlapped with the reproductive period of *C. briggsae*. Comparing DEGs between these four time points (i.e., day 1, day 3, day 6, and day 9) allowed us to determine gene expression changes during and after reproduction.

In the *C. briggsae* transcriptome, we found that some large-scale patterns were noticeable in the DEGs during various time points sampled between day 1 and 9. When examined without any log2 fold cutoff, roughly the same proportion of genes were up- and down-regulated at all measured time intervals (Supplemental File 1; Supplemental Table 2). However, the size of expression changes was less neutral, after application of a cutoff at 1.0 log2 fold (Table 1). A comparison of day 1 to day 3 (referred to as ‘day 1-3’) revealed that the majority of DEGs were downregulated by day 3 (Figure 2; Table 1). A similar trend was observed in day 1- to-day 6 comparison (day 1-6), although when compared to day 3 (i.e., day 3-6), where very few genes were differentially expressed, more genes were upregulated instead. This shift towards upregulation continued in the later time point of day 6 compared to day 9 (day 6-9). The strong bias towards downregulation by day 3, at roughly the mid-reproductive stage and upregulation afterwards may indicate a shift in transcriptome as the animals begin transition into a post-reproductive physiological state.

**FIGURE 2.**
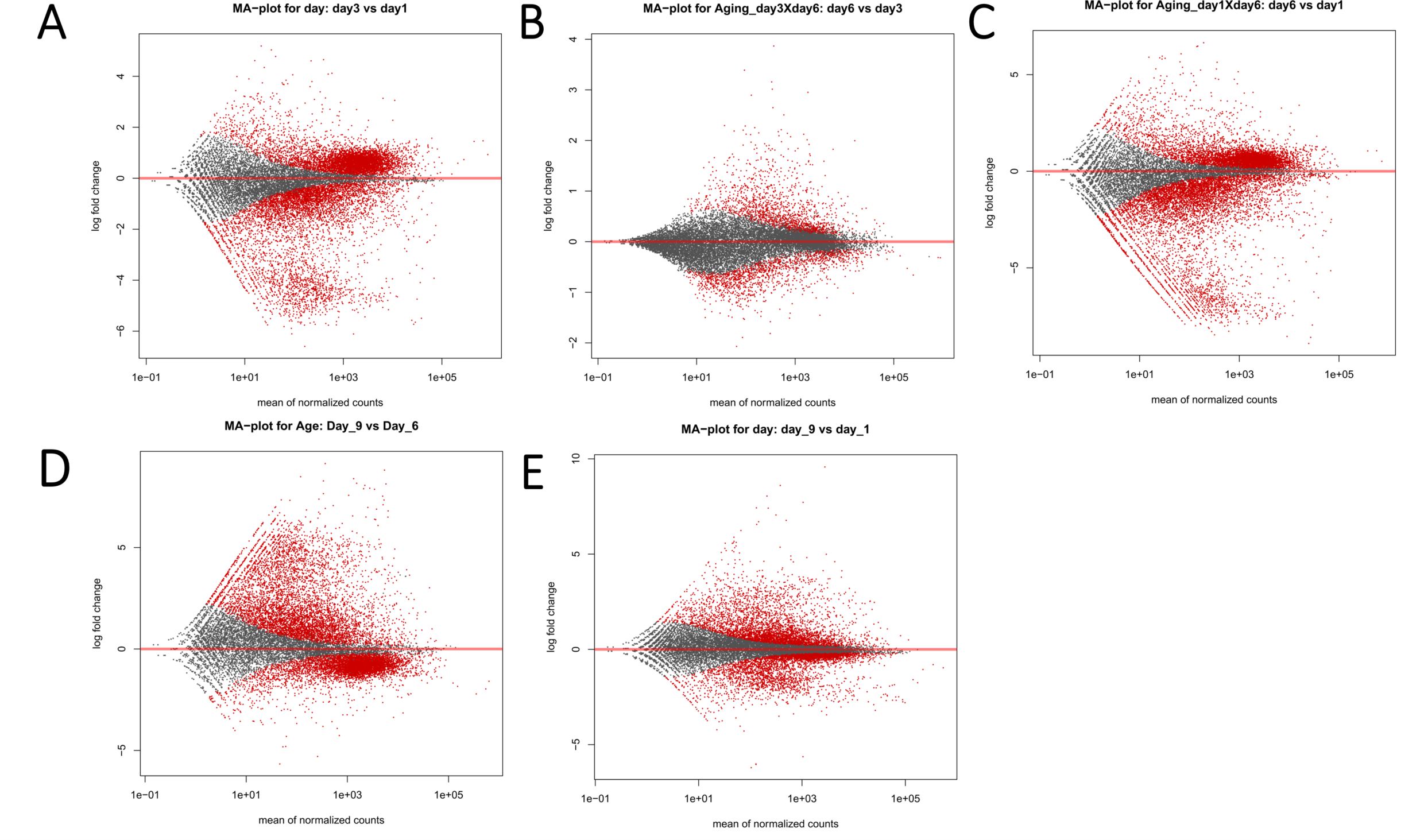
MA plots of differentially expressed genes from day 1, 3, 6, 9. **A-E**. Differentially expressed genes between Day 1-3 (A), 3-6 (B), 1-6. (C) 6-9. (D) and 1-9 (E). Red dots indicate significantly regulated DEGs, grey dots are not significantly regulated.

**Table 1.**
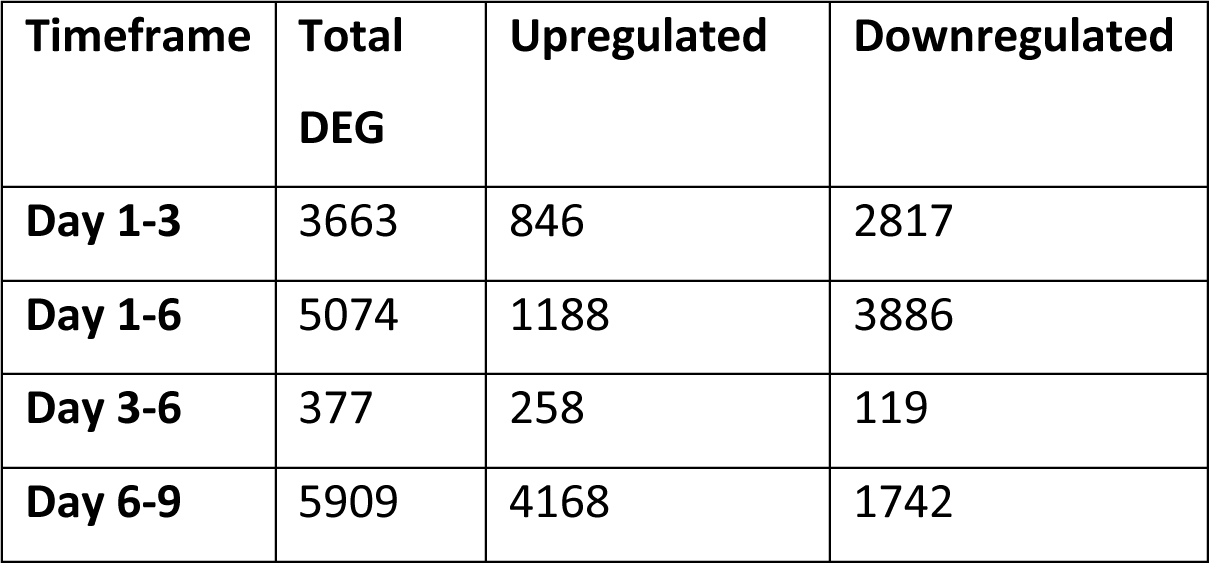
Age-based differentially expressed *C. briggsae* genes. Numbers of DE genes (p<0.01) obtained from pairwise comparison of RNA-sequencing on differently aged synchronized *C. briggsae* populations. The table contains numbers of genes after applying a minimum 2-fold expression change cutoff (log2fold expression +/−1).

### Analysis of DE genes during reproductive and post-reproductive stages

As discussed above, *C. briggsae* hermaphrodites are reproductively active untill day 6. Hence, day 3 and 6 can be compared to day 1 to examine gene expression changes within the reproductive period. Comparison with the day 9 transcriptome should reflect post-reproductive expression changes. The genes in the comparisons of day 1-3 and day 1-6 are strongly overlapping (3,117 of 3,663, i.e., 85%; Figure 3a,b). All but 4 of the overlapping genes maintain the same directionality, further confirming that genes that are activated during reproduction maintain their expression trends untill day 6.

**FIGURE 3.**
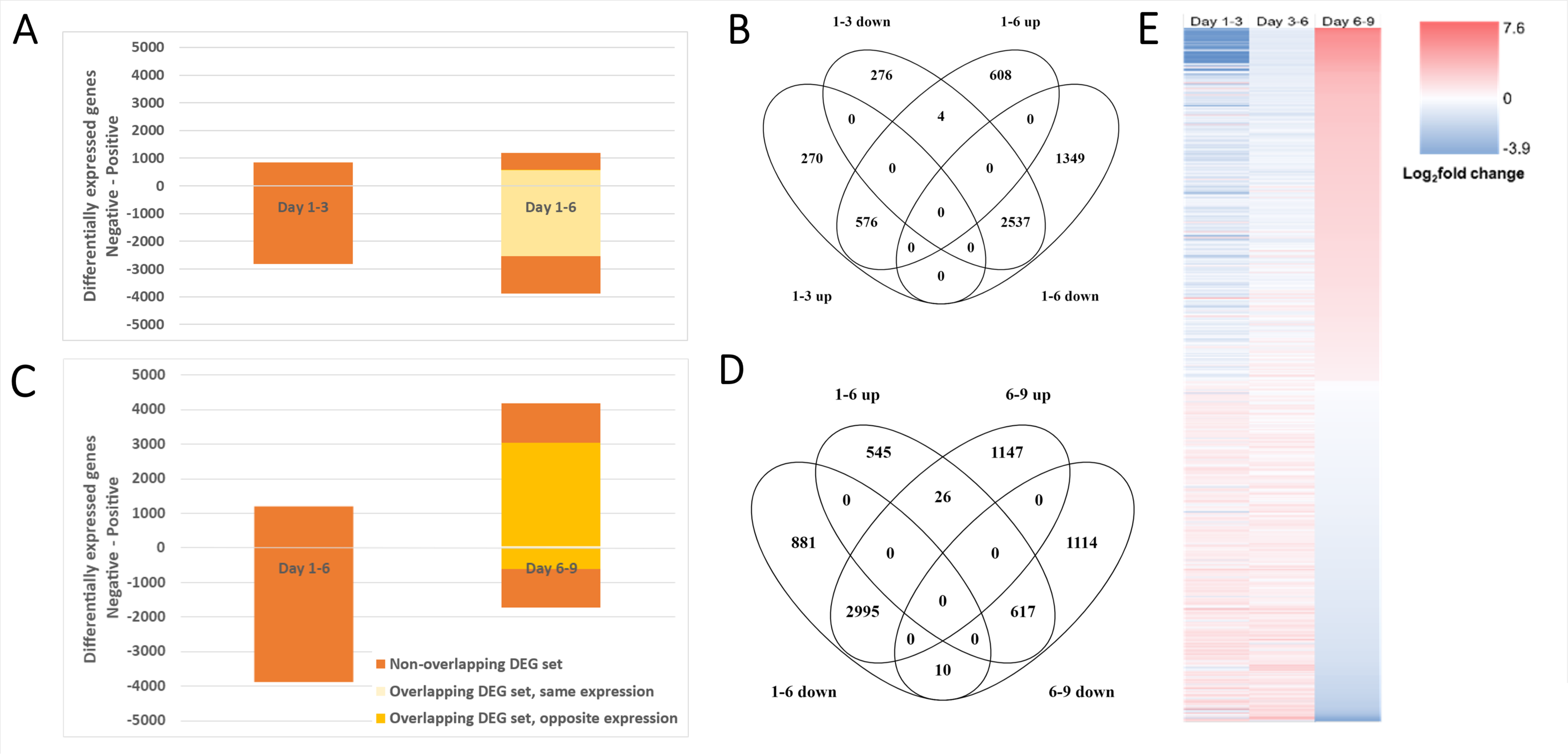
Comparison of gene expression direction changes with time. **A,B.** Bar graphs of day 1-3 vs 1-6 (A) and 1-6 vs 6-9 (B). Bars on the right are labelled to differentiate between genes not included in the left timepoint set (red), genes included in the left set and having the same expression change direction, i.e. both increasing or decreasing (light yellow), and genes included in the left and having an opposite expression change direction (dark yellow). Inclusion criteria were relative log2fold expression +/−1, p<0.01. **C,D.** Venn diagrams of 1-3 and 1-6 DEGs (C), 1-6 and 6-9 DEGs (D). Inclusion criteria were relative log2fold expression +/−1, p<0.01. **E.** Heatmap of all 687 *C. briggsae* genes with significant expression changes at each pairwise measured interval between adult day 1, 3, 6, and 9. p<0.01.

Further comparison of day 1-6 revealed 1957 genes that are not part of the 1-3 set (Figure 3a,b). Most of these (89%, 1736) changed their expression very slowly such that they were not part of day 1-3 and day 3-6 sets (Supplemental File 1). While the biological significance of this slow increase trend remains to be determined, it would be unlikely that these genes are involved in initiation of reproduction but might play a role in its maintenance. Interestingly, a comparison of DEGs between day 3 and 6 gave significantly fewer genes, specifically 377 (Day 3-6 set; 258 up and 119 down). This suggests that animals remain in a comparatively transcriptionally stable state from day 3 to 6. Filtering for significant changes at each interval showcases how gene expression radically shifts after the reproductive phase, with 687 genes in total (Figure 3e).

Comparing gene expression changes between day 1-6 and day 6-9 sets revealed an interesting trend of expression reversal (Figure 3c,d). We found that 52% (617 of 1188) of upregulated genes reversed the direction of their expression, *idem* for 77% (2295 of 3886) of the downregulated genes. Up-invert genes (i.e., genes upregulated during reproduction and downregulated post-reproduction) had a close range of expression change, with a mean log2fold expression change of 0.80 from day 1 to 6 (figure 4a). By day 9 these genes returned to levels comparable to day 1 (mean −0.16), suggesting that they may not play a major role beyond reproduction. The expression changes of down-invert genes (i.e., genes that were downregulated during reproduction and upregulated post-reproductively) were much larger and have a greater spread on day 6 with a mean of –2.87 with percentiles of –4.63 and –1.16, and on day 9 a mean of 0.19 with percentiles of –1.00 and 0.47. We further analyzed the down-invert set (blue), which revealed that 1121 genes remained downregulated by day 9 (blue), 1620 returned to levels not significantly different from day 1, and 1135 genes were greatly increased, beyond day 1 levels (purple) (Figure 4b). While these expression differences may indicate how different sets of genes might function, we suggest that down-invert genes may be involved in the physiological processes during the post-reproductive period of life.

**FIGURE 4.**
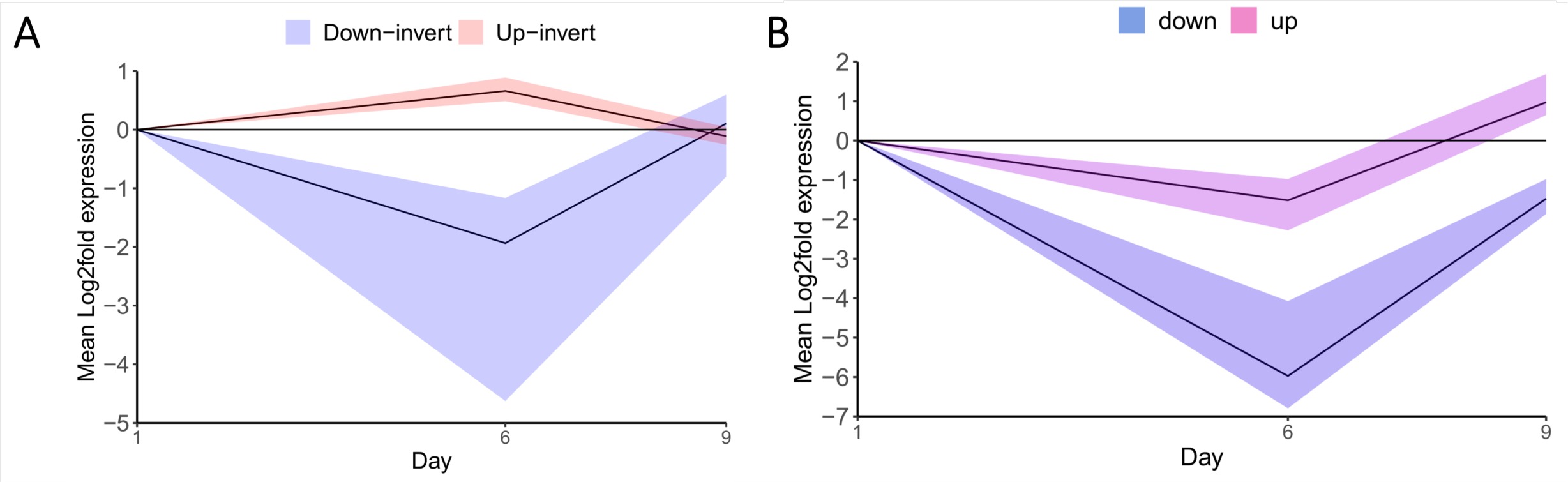
Gene sets with inverted expression post-reproduction. **A.** Median with 75 percentile (shaded area) genes upregulated (red) and downregulated (light blue) on day 6 compared to day 1 of adulthood, with expression inverted to downregulated and upregulated by day 9, respectively. **B.** Median and down-inverted gene set upper and lower 25 percents (shaded area) in violet and dark blue respectively, with shaded areas up to minimum and maximum expression values.

### Enrichment analysis of processes and pathways that correlate with reproductive shift

To better understand changes in physiological processes that occur as gene expression shifts from the reproductive (1-6) to post-reproductive (6-9) stage of life, GO analysis was performed for *C. briggsae* genes and their *C. elegans* orthologsm, along with KEGG analysis on the latter. While both approaches produced largely similar results, *C. elegans* ortholog analyses provided a larger set of annotation terms and a higher coverage than *C. briggsae*. Altogether seven major categories were identified each consisting of related GO biological process terms and KEGG pathways: matrisome, muscle development and function, DNA damage repair, stress and immune response, fatty acid metabolism, catabolism (non-lipid), and regulation of transcription. Figure 5 shows general expression trends of associated genes. The Supplemental file 2 contains a full list of the grouped GO and KEGG terms with enrichment scores.

**Figure 5:**
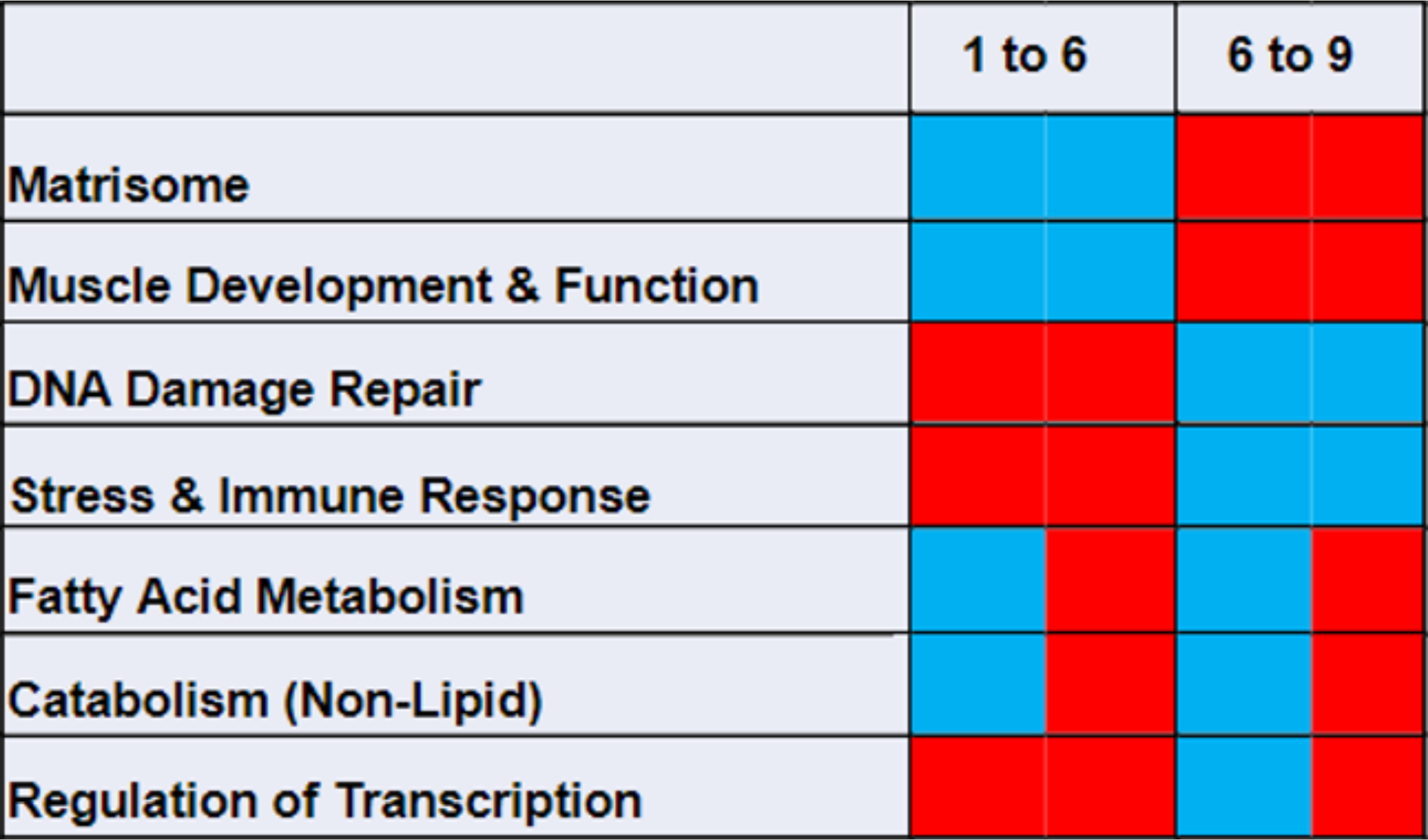
*C. briggsae* longitudinal transcriptome GO and KEGG term categories. GO term expression change direction between day 1, 6 and 9 of adulthood – statistically significant, log2fold cutoff +/− genes list. Positive enrichment of upregulated genes and negative enrichment of downregulated genes both refer to related processes becoming more active (red). Conversely, positive enrichment of downregulated genes and negative enrichment of upregulated genes refer to related processes becoming less active (blue). Boxes with both colours indicate a mixture of GO terms that are divided between both categories. For details on GO and KEGG annotation analysis, see Methods.

The overview of the major categories revealed three broad expression trends during reproductive to post-reproductive period shift, i.e., gene expression initially going down and then up (matrisome and muscle development & function), going up and then down (DNA repair and stress & immune response), and a mixed direction change (fatty acid metabolism, lipid and non-lipid catabolism and transcription & transcription factors). In general, both GO terms and sets of associated genes showed similar enrichment trends. The above categories and GO terms are briefly discussed below, with additional descriptions available in the supplemental text.

We observed a significant enrichment of sets of matrisome genes that were upregulated post-reproduction. A comparison of DEGs between the reproductive (day 1-6) and post-reproductive (day 6-9) periods revealed genes linked to ‘basement membrane’ and categories such as ‘extracellular matrix organization’ and ‘collagen-containing extracellular matrix’. These were significantly downregulated during the reproductive phase (day 1-6) but were upregulated post-reproductively (Supplemental Figure S1a). The set of genes includes laminins, transcription factor *fos-1*, collagens (*Cbr-emb-9* and *Cbr-let-2*, *Cbr-cle-1*), papilin *Cbr-mig-6*, nidogen *Cbr-nid-1*, SPARC, and the proteoglycan *Cbr-unc-52*. Since collagens were strongly represented among the genes that show upregulation after reproduction, we analyzed these in further detail through comparison with previous findings. Of the 181 *C. elegans* cuticular collagens (Teuscher et al., 2019), 149 were present in the *C. briggsae* genome as orthologs (Supplemental File 3, Supplemental Figure S1b). Of these, 77 genes also underwent the down-invert pattern: initial downregulation, then revert to upregulation. The set included the 4 collagens with human orthologs (*Cbr-cle-1, Cbr-col-99, Cbr-emb-9* and *Cbr-let-2*).

Like the matrisome pattern of expression, we observed gene expression profiles for several terms related to muscle development and function (Figure 5). Specifically, there was reproductive period downregulation and post-reproductive upregulation for genes involved in ‘regulation of striated muscle cell differentiation’, ‘muscle cell cellular homeostasis, ‘muscle system process’ and ‘muscle contraction’, ‘myofilament’, ‘sarcomere’, ‘contractile fiber’ and ‘striated muscle dense body’. Some examples included *Cbr-mup-2*, *Cbr-pat-10* and tropomyosin *Cbr-lev-11*, which together regulate myosin-actin interactions when activated by calcium influx (Myers et al., 1996), and members of gene families such as troponins and myosins. Further, there were twitchin *Cbr-unc-22* and titin *Cbr-ttn-1*, sarcomere proteins that regulate muscles to adapt to changing physical demands as usage fluctuates through life (Matsunaga et al., 2015). (Schorr et al., 2023) recently published a comprehensive list of *C. elegans* body muscle expressed genes. Although these were in mixed stage animals, we found that *C. briggsae* orthologs of body-muscle specific genes *Cbr-unc-54*, *Cbr-mlc-2.2*, *Cbr-unc-27* and *Cbr-lev-11* all fit in the down-invert pattern, and a majority of muscle-specific genes with RNA-binding domains as well (Supplemental file S4). Several genes were related to an interaction between muscle and the basement membrane, including perlecan *Cbr-unc-52*, talin *Cbr-tln-1* and the integrins *Cbr-pat-2* and *Cbr-pat-3*, all of which are required to maintain muscle motor functions (Etheridge et al., 2015).

Unlike the above two categories, GO terms related to DNA damage repair were enriched for genes that were post-reproductively downregulated. The pattern of expression changes was unique in that the upregulated genes in day 1-3 set were enriched for terms involved in DNA damage repair or DNA replication but not in day 3-6 and day 1-6 sets, indicating that between day 1 and 3, these processes are most active. Early reproductive phase enrichment included ‘DNA repair’ and ‘double-strand break repair via break-induced replication’, with several MCM (minichromosome maintenance) complex components including *mcm* family members (*Cbr-mcm-2*, *Cbr-mcm-3*, *Cbr-mcm-4*, *Cbr-mcm-5*, and *Cbr-mcm-6*), *Cbr-lem-3* and DNA repair protein *Cbr-rfs-1/RAD51*. There were also many terms involved in DNA replication, some of which overlapped with DNA repair.

The stress and immune response genes showed a largely similar trend as the DNA damage repair set, except that the expression trend was significantly upregulated in both day 1-3 and day 1-6 sets, followed by downregulation post-reproductively. This trend suggests that stress and immune response processes are active throughout the entire reproductive phase. Representative genes and gene families that showed reproductive shift include three caenacins (*Cbr-cnc-4, Cbr-cnc-5.2, Cbr-cnc-11*), seven lysozyme-like protein genes (*lys* & *ilys* family) of which *C.elegans* orthologs are involved in immune response against bacterial infection (Nicholas and Hodgkin, 2004) and systemic stress signaling *Cbr-sysm-1*, ortholog of *C. elegans sysm-1*, a target of *pmk-1* inducing apoptosis in response to environmental stressors (Soltanmohammadi et al., 2022). Other examples included the *hsp-16.41* ortholog *CBG04607* and four *hsp-16.2* (HSP16 family) orthologs that displayed the reproductive shift.

KEGG pathway annotation for day 1-6 DEG was enriched for the lysosome pathway including three *cpr* genes (*Cbr-cpr-1*, *Cbr-cpr-3*, *Cbr-cpr-4*). In *C. elegans*, *cpr* class genes, together with caenacins *cnc-1* and *cnc-2*, have been shown to increase expression under influence of CLEC-47 overexpression, a signaling protein that activates the immune response (Pan et al., 2021). The upregulation of immune and stress response genes during the reproductive phase and subsequent downregulation is consistent with evolutionary pressure to maximize survival during the reproductive period and reproductive success.

As mentioned above, fatty acid metabolism consisted of GO terms that displayed a mix of expression trends. While some GO terms were enriched among genes that became down-then upregulated, others represented processes that showed an opposite trend. Within this group fits the set of terms involving metabolism and catabolism of fatty acids. The downregulated genes during the reproductive phase were enriched for four fatty acid elongation GO terms, each containing the same seven long chain fatty acid elongation genes (*Cbr-elo-1*, *Cbr-elo-2*, *Cbr-elo-4*, *Cbr-elo-5*, *Cbr-elo-7*, *Cbr-elo-8*, and *Cbr-elo-9*). Post-reproductively, annotation was enriched for fatty acid and lipid metabolism terms. The genes that invert expression upwards after reproduction were enriched for terms such as ‘lipid metabolic process’, ‘lipid catabolic process’ and ‘fatty acid catabolic process’. These included 20 lipase genes with seven *lips*-family lipase genes and *Cbr-lipl-7*, as well as fatty acid hydrolases *Cbr-faah-1*, *Cbr-faah-3*, and three acyl COA dehydrogenases (*CBG18107*, *CBG15945*, *CBG02501*).

Among the post-reproductively downregulated genes, we found enrichment for several terms related to lipid metabolism. The terms included ‘negative regulation of lipid metabolic process’, ‘lipid metabolic process’ and ‘lipid biosynthetic process’, including *Cbr-elo-3*, as well as *Cbr-faah-2*, *Cbr-faah-4*, *Cbr-faah-5*, *Cbr-lipl-1*, *Cbr-lipl-2*, and *CBG10449*, which are all members of the same gene classes in the reproductively downregulated set but not the same genes. Under ‘lipid localization’ we found all five *C. briggsae* vitellogenin genes (*CBG16767*, *Cbr-vit-2*, *CBG14203*, *CBG14234*, *Cbr-vit-6*). Vitellogenins are involved in the balance between reproductive success and lifespan as they traffic lipids from the intestine to eggs (Kern et al., 2023; Ranawade et al., 2018).

In agreement with GO annotation, when analyzing the KEGG pathways of *C. elegans* 1- to-1 orthologs, ‘fatty acid metabolism’ and ‘fatty acid biosynthesis’ were enriched among both up and downregulated genes during the reproductive period. However, for the post-reproductive period, enrichment was observed only among upregulated genes. Furthermore, we found that the entire set of post-reproductive DE genes were enriched for the ‘mTOR signaling pathway’ that is involved in lipid metabolism (Lamming and Sabatini, 2013). *mom-2* (*Wnt*) and *dsh-2* were down, reducing inhibition of GSK3β and thereby reducing inhibition of mTORC1. Similarly, *strd-1* (*STRAD*) was downregulated, impairing phosphorylation of *aak-1/aak-2* (*AMPK*). Affecting *mTORC2*, phosphorylation factor *aap-1* (*PI3K*) was also among the downregulated genes.

Besides lipid catabolism we noted that many other catabolic processes also were affected by the reproductive period expression shift, and that the genes involved rarely overlapped with the lipid catabolism sets, indicating that these were distinct processes. The GO terms ‘small molecule catabolic process’, ‘amino acid catabolic process’ and ‘carboxylic acid catabolic process’ were all enriched among down-invert genes. The analysis of various affected terms suggested an overall downregulation of proteasome-mediated catabolism in the post-reproductive period, which is consistent with previous studies showing cellular machinery becoming less efficient in dealing with stress with age.

The final category entails regulation of gene transcription. Genes such as transcription factors are important to consider in transcriptomic analysis, given that changes in their expression could have profound downstream consequences. Finding germline-associated transcription factors was especially relevant considering the reproductive shift expression, and a set are described in detail further on. Our analysis revealed that on day 3, upregulated genes were enriched for several terms related to general regulation of transcription, including genes *Cbr-egl-18*, *Cbr-glp-1*, and *Cbr-lin-35*. However, interestingly, by day 6, no GO term showed significant enrichment among upregulated genes. When examining the post-reproductive period, the expression trend changed to enrichment of processes for genes both involved in specifically positive regulation as well as negative regulation of transcription (Supplemental file 2).

To further investigate the genes affecting transcription, we analyzed the expression of the RNA polymerase II core complex members in our dataset. The analysis showed that during reproduction, expression of these genes either remained unchanged or increased significantly whereas post-reproductively, all of them were decreased (Supplemental file 5). A similar expression analysis of corresponding *C. elegans* genes using published datasets (Schmeisser et al., 2013) revealed increases during reproduction for only 3 core complex genes, and no significant downregulation post-reproductively. Thus, the effect appears to be specific to *C. briggsae*.

It was recently reported that RNA processing fidelity is affected by aging (Ham et al., 2022). RNA processing becomes less efficient with age, with overall RNA quality decreasing as various non-exon protein-coding and non-coding RNA categories become more prevalent. We analyzed the RNA processing related terms in *C. briggsae* (Supplemental file 2) and found these to be enriched among up-invert genes, consistent with the process becoming less active in post-reproductive age.

### Comparison of gene expression trends between *C. elegans* and *C. briggsae*

We compared our longitudinal *C. briggsae* transcriptome with published *C. elegans* datasets that feature reproductive and post-reproductive timepoints (Schmeisser et al 2013). The analysis of differentially expressed genes revealed a similarity in the reproductive period downregulation bias (Figure 6). Specifically, there were 2.5 times more downregulated genes compared to those that were upregulated (Supplemental table 3).

**Figure 6:**
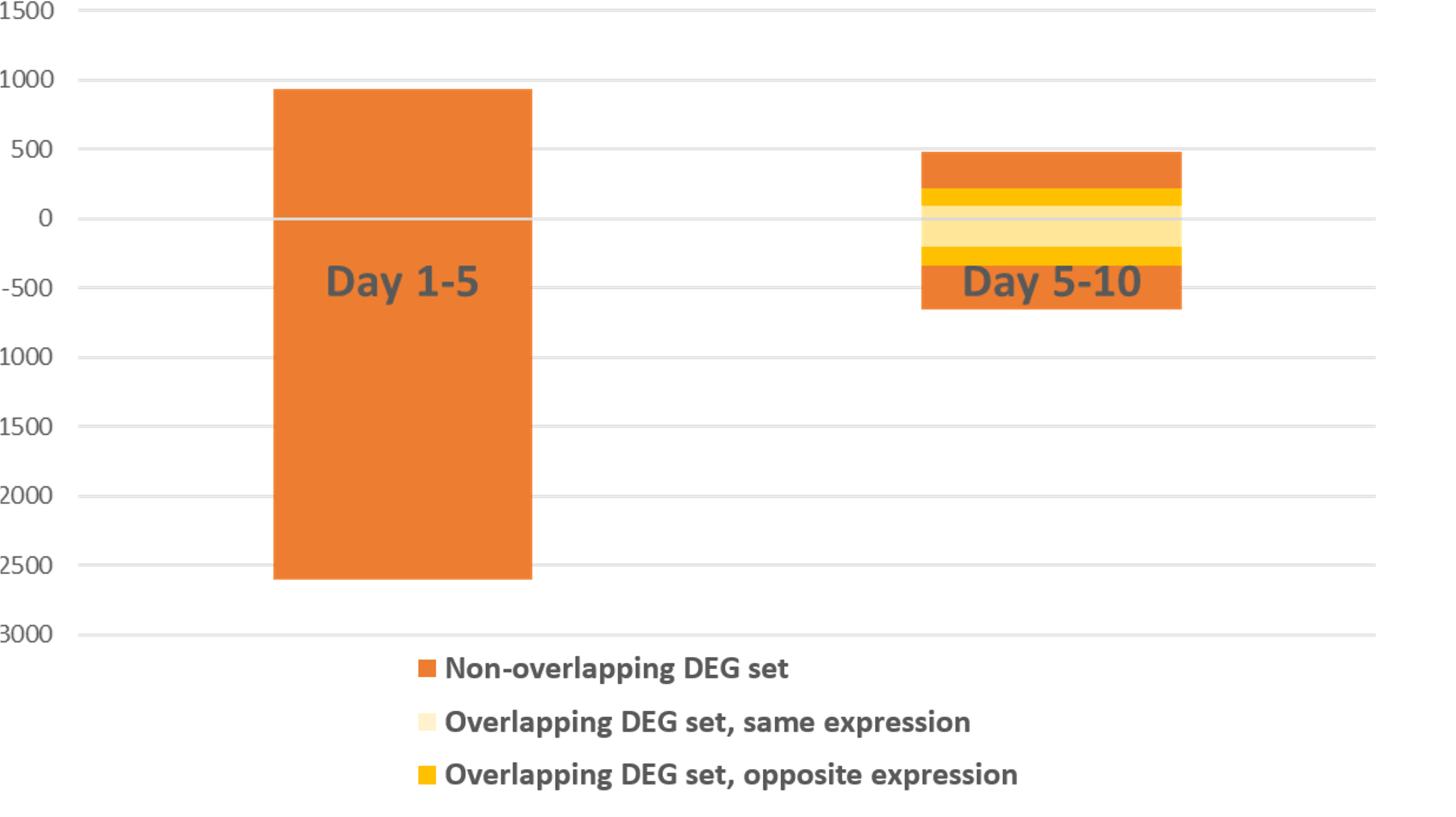
Comparison of gene expression direction changes with time in *C. elegans*. Bar graph of day 1-5 vs 5-10 DEGs in *C. elegans*. Bars on the right are labelled to differentiate between genes not included in the left timepoint set (red), genes included in the left set and having the same expression change direction, i.e. both increasing or decreasing (light yellow), and genes included in the left set and having an opposite expression change direction (dark yellow). Transcriptome DEG data used from (Schmeisser et al., 2013). Inclusion criteria were relative log2fold expression +/−1, p<0.01.

Interestingly, *C. elegans* did not display a reproductive shift as stark as we found in *C. briggsae*: during the post-reproductive period the downregulation bias disappeared and no major upregulation bias was seen. However, the analysis of GO terms from the Golden et al. study (Golden et al., 2008) revealed the presence of several of the annotation categories that were observed earlier in the *C. briggsae* transcriptome reproductive shift.

The matrisome terms were found among downregulated genes during reproduction, similar to *C. briggsae*, but post-reproductively there was no enrichment among upregulated genes. We took a closer look at a set of 20 *C. elegans* collagens having reduced expression until day 5 of adulthood followed by a slow increase with age (Golden et al., 2008). Of the 13 orthologs present in *C. briggsae*, we found a conserved, down-invert expression pattern for 11 of these: *Cbr-rol-6*, *Cbr-col-34*, *Cbr-col-49*, *Cbr-col-73*, *Cbr-col-122*, *Cbr-col-184*, *Cbr-dpy-5*, *Cbr-dpy-13*, *CBG18259*, *CBG23158*, and *CBG24927*. Only *Cbr-col-101* and *Cbr-col-158* show species-specific expression patterns, with the former decreasing expression by day 9 and the latter increasing on both days 6 and 9.

Muscle development related terms were enriched among down-invert genes, but there was no significant enrichment for muscle function terms. This is different from *C. briggsae*, where a down-invert pattern was seen for both terms suggesting that collectively muscle genes are regulated differently. Orthologs of the *C. briggsae* upregulated muscle genes were not upregulated in *C. elegans* when compared to the Schmeisser et al. dataset (Schmeisser et al., 2013). However, a comparison of our *C. briggsae* reproductive shift genes to orthologs of muscle-tissue expressed genes in *C. elegans* adults (Kaletsky et al., 2018) yielded an overlap of 97/325 with up-invert genes (29.8%) and 395/2,235 down-invert genes (17.7%) (Supplemental file 6). This suggests that 19.2% of all muscle-related DEGs in our dataset have *C. elegans* orthologs that are enriched in muscle tissue, and a majority of these *C. briggsae* muscle genes represent the whole-transcriptome pattern of downregulation during reproduction.

### Germline genes analysis

We reasoned that genes that are expressed in reproduction and show a reproductive shift would correlate with reproductive processes. To investigate this, we examined expression data of *C. elegans* germline genes (Reinke et al., 2004). Our analysis showed that germline genes were highly enriched among the *C. briggsae* genes that were upregulated during reproduction and downregulated post-reproductively (103 of 222 genes, hypergeometric test p<0.001) (Supplemental figure S2, Supplemental file 7). Thus, a significant proportion of up-invert genes are involved in reproduction. The overlapping genes are linked to a broad range of processes, including oocyte development and maturation, embryogenesis, and embryonic development. The up-invert germline set includes *Cbr-puf-8*, for which the *C. elegans* ortholog is known to inhibit the overproliferation of germline stem cells in the germline allowing for stem cells to enter meiosis and become oocytes (Racher and Hansen, 2012).

Reproduction of hermaphroditic nematodes involves many genes, which can be split up into components such as oogenesis, spermatogenesis, and intrinsic germline related genes (Reinke et al., 2004). Work in *C. elegans* has shown that transcription factors *lin-35*, *efl-1*, and *dpl-1* are necessary for proliferation of germ cells and affect expression of several downstream genes, although only *efl-1* and *dpl-1* are essential for oogenesis (Chi and Reinke, 2006). Our expression analysis in both species revealed that in *C. elegans,* all three genes are upregulated during the reproductive period, however post-reproductive downregulation is observed only in *C. briggsae*. The analysis of downstream genes affected by these three factors showed that their genetic networks were largely conserved, with some species-specific differences (Supplemental File 8). Whereas the expression changes of *lin-35* regulated genes were evenly split in *C. elegans*, most were upregulated in *C. briggsae*.

Another transcription factor, *spe-44*, promotes spermatogenesis (Ragle et al., 2022) and may also have a conserved role in *C. briggsae* (Kulkarni et al., 2012). Interestingly, we found that the *Cbr-spe-44* expression has an up-invert pattern, unlike its *C. elegans* ortholog that remains unchanged through adulthood. Overall, these expression patterns of transcription factors and downstream genes suggest specific differences between *C. elegans* and *C. briggsae* reproductive gene expression, which might help explain the exclusive presence of the reproductive shift effect in *C. briggsae*.

### Manipulating the reproductive state of animals affects gene expression

If the reproductive shift is related to the reproductive state of animals, then manipulating that state should affect gene expression. To test this, we selected a random set of three up-invert (*Cbr-clec-266, Cbr-eor-1*, and *Cbr-unc-71*) and three down-invert genes (*Cbr-col-129, Cbr-col-139*, and *Cbr-mpz-4*) (Supplemental table 4). These genes represented germline, collagen, immune response, and transcriptional regulation processes. The qPCR analysis was performed in animals that were either sterile, using *Cbr-glp-4(v273)* (Velayudhan and Ellis, 2022) or had reproduction extended by mating. The up-invert and down-invert genes in the *Cbr-glp-4(v273)* mutant condition were expected to not undergo the reproductive shift. In the other condition, day 9 mated animals were expected to be similar in expression to day 3 controls, and dissimilar to day 9 unmated animals. We found that in some cases experimental manipulation affected gene expression (Supplemental Figure S3, Supplemental Table S4), however further conclusions could not be drawn due to variance among replicates. In the case of *Cbr-mpz-4,* we observed expected expression change, i.e., upregulated after mating on both day 3 and 9 compared to unmated animals (Supplemental Figure S3b) but post-reproductive day 9 unmated animals did not differ from day 3 unmated controls. Several factors are likely to affect our data and observed variabilities, for example, *Cbr-glp-4(v273)* mutants requiring high temperature to induce sterility, frequency and duration of male mating, and the secondary effect of male sperms on hermaphrodites (Scharf et al., 2021; Velayudhan and Ellis, 2022). These experimental manipulations are known to alter gene expression beyond affecting just the reproductive state (Devanapally et al., 2021; Golden and Riddle, 1984; Klosin et al., 2017; Wang et al., 2021). Future experiments should be performed with a larger set of genes along with additional experimental conditions such as mating with sterile males or using gonad ablations to induce sterility, in order to draw firm conclusions about the biological effects of reproductive shift. Overall, while results of experimental manipulations did not allow us to affect the reproductive shift in *C. briggsae* in a predictable manner, more work is needed to investigate the effect of reproduction on gene expression changes in a fulsome manner.

## DISCUSSION

We performed transcriptome analysis on several timepoints spanning the reproductive and post-reproductive periods of *C. briggsae*. The results revealed a reproductive shift, where the expression of a majority of the DE genes changed in an opposite direction post-reproductively. We categorized the higher-order physiological processes affected by either up-invert or down-invert expression patterns through annotation analysis, examined alterations in germline genes, transcription factors, and transcriptional networks, and performed a comparative study in *C. elegans*. We also experimentally confirmed expression changes for several genes in *C. briggsae* by manipulating the reproductive state of animals.

Our transcriptomic study of *C. briggsae* hermaphrodites showed several interesting patterns of gene expression, with the reproductive shift effect chiefly among them. We found that large portions of reproductive period DEG’s undergo a strong change in expression trend with the cessation of reproduction. Simultaneously, very few genes showed further increase or decrease in the same direction. We suggest that the cessation of reproduction plays a major role in the observed gene expression pattern, besides the process of aging itself. The larger expression change seen upon entry into the post-reproductive period for *C. briggsae* is remarkable. Interestingly, while the gene expression pattern during the reproductive phase in *C. elegans* was found to be similar to *C. briggsae*, the post-reproductive change showed differences. It should be noted that *C. briggsae* has many notable dissimilarities from *C. elegans*, with developmental, genetic and genomic changes underlying phenotypic differences in temperature and stress sensitivity among other facets (Gupta et al., 2007; Jhaveri et al., 2023). Although both species reproduce hermaphroditically, this was not true for the shared common ancestor (Kiontke et al., 2004). Our findings of transcriptomic differences further highlight the distinct physiologies of these two species.

Through annotation analysis, we associated several major processes with gene expression changes seen in the reproductive shift. The most consistent patterns we found were down-invert regulation for matrisome and muscle related genes, and up-invert regulation of DNA damage repair, stress and immune response related genes. Collagen is a major component of the matrisome, and expression of some collagen genes increases with age (Halaschek-Wiener et al., 2005). Phenotypically, cuticle thickness also increases, which has been suggested to be the result from more lax expression regulation after the reproductive phase (Herndon et al., 2002). A reduced evolutionary pressure in post-reproductive gene expression has also been theorized, since gene expression patterns will no longer be able to confer a direct fitness benefit to offspring (Rose, 1991), that is, assuming the hermaphrodite does not receive new sperm from a mating event. Our analysis of collagens in *C. elegans* revealed a similar expression trend as in *C. briggsae*. Intertwined with the progression from reproductive to post-reproductive stage are also age-related effects on expression of certain collagen and cuticle-related genes that were reported to decline in older adults (Budovskaya et al., 2008; Ewald et al., 2015; Palani et al., 2023). Cuticular collagens have been found to confer resistance to pathogen infection and oxidative stress (Ewald et al., 2015; Sellegounder et al., 2019), which wane as animals age (Lopez-Otin et al., 2013). Either a full relaxation, or a partial relaxation of selection pressure after the reproductive period would fit the data presented here. However, the chance of a post-reproductively aged, sperm-depleted hermaphrodite encountering a male and restarting reproduction is non-zero, which leads to the assumption that some amount of selection pressure remains towards maintaining fitness in post-reproductively aged *C. briggsae*. Alternatively, and somewhat more likely based on the specific enrichments for collagens and the matrisome, these genes remain or become important for the maintenance of reproductive fitness after the self-reproductive period. This would make collagens and the matrisome viable targets for intervention towards the maintenance of post-reproductive health.

Similar to the matrisome genes, muscle-related genes in *C. briggsae* showed increased expression post-reproductively. Correct muscle function is dependent on anchoring to other cells and matrisome components and therefore would be expected to show analogous gene expression changes. Interestingly, in *C. elegans*, expression of muscle-related genes also decreased during the reproductive period but did not increase afterward. Furthermore, it was reported earlier that *C. elegans* muscle-specific gene transcripts, particularly those involved in muscle contraction, progressively and consistently became downregulated until day 7 of adulthood (Mergoud Dit Lamarche et al., 2018). Another recent study reported that gene expression in the muscle tissue of *C. elegans* did not increase with age (Roux et al., 2023). Thus, while in *C. briggsae* there is a pattern of post-reproductive upregulation for muscle-related genes, in *C. elegans* there is no such clear pattern which is an interesting difference. Muscle-related genes have been found to show a mix of transcriptional pattern across organisms. In *Drosophila melanogaster*, muscle function and development genes are downregulated in muscle tissue measured in the thorax of older animals, while proapoptotic factors become upregulated in the same tissues coinciding with muscle breakdown (Bordet et al., 2021; Girardot et al., 2006). Zhan et al. saw that at post-reproductive age, reproductive and muscle tissues had similar expression profiles, with the downregulated genes enriched for muscle contraction, muscle development and myogenesis (Zhan et al., 2007). However, interestingly, studies in rats and humans have reported gene expression in skeletal muscle tissue, including several genes involved in muscle development, skewed towards increasing with age (Shavlakadze et al., 2019; Tumasian et al., 2021). It is possible that the different ways of measuring, i.e. whole-animal and tissue-specific transcriptomes, cause the different expression patterns associated with muscle function with age. It may be that upregulation is related to the repair and maintenance of muscle tissue as animals transition to the post-reproductive phase of life. More work is needed to understand the biological significance of the expression trend and whether this would translate into more protein production and their relationship to muscle function.

Cellular stress, such as mitochondrial stress and increased reactive oxygen species generation is thought to contribute to the physiological decline that occurs with aging (Shpilka and Haynes, 2018). Our analysis showed that on a whole-genome level, stress response processes decline in older, post-reproductive animals. Various individual stress response genes varied: In *C. elegans*, heat-shock proteins as a group showed no consistent trend, and neither did we find enrichment for oxidative stress factors. It is unclear whether changes in stress genes are associated with the reproductive state or aging, although *hsp-16.2* gene expression has been found to correlate with and predict lifespan (Mendenhall et al., 2012). Post-reproductive decline of stress response genes has been found in *C. elegans*, albeit steadily, not sharply after reproduction (Golden et al., 2008; Walker et al., 2001).

DNA repair-related genes were upregulated in *C. briggsae* during the early period of reproduction (day 1 to day 3) with a smaller inferred decline until day 6 and a significant decline during the post-reproductive phase. There was no enrichment for relevant terms in our *C. elegans* analysis. Another study suggested that increases in chromatin condensation and DNA damage in older *C. elegans* lead to decreasing gene expression fidelity, since the authors observed increases in transcription factor expression in day 9 adults and reasoned it to be a potential compensatory mechanism (Golden et al., 2008). More generally, including in humans, DNA repair is known to decrease with age, both on the transcriptional level as well as in effectivity (Li et al., 2016; Lidzbarsky et al., 2018). The decline of DNA repair with age measured as genomic instability and telomere attrition is common among many species and is seen as an important factor limiting lifespan, although more programmatic theories of aging should also be considered (de Magalhaes and Church, 2005; Lopez-Otin et al., 2013; Medvedev, 1990).

Among other processes, lipid metabolism and catabolism related genes appeared to vary in expressivity throughout the stages measured. Expression of vitellogenin genes matches with the initial high energy investment in egg production. This first decreases moderately while egg production tapers down, and after oocyte stores are depleted, continues to traffic lipids for some time to produce ventable yolk, to support offspring (Kern and Gems, 2022; Kern et al., 2021). Finally, the analysis of germline transcription factors *Cbr-lin-35*, *Cbr-efl-1*, *Cbr-dpl-1*, and *Cbr-spe-44* showed an up-invert expression in *C. briggsae* but no such pattern in *C. elegans*. Moreover, the expression of genes acting downstream of these factors showed that the species-specific reproductive shift effect has a large effect on reproduction related factors in *C. briggsae*.

It is important to point out that transcriptomic changes during the reproductive phase not only reflect processes in mothers but also developing oocytes in the uterus. Many longitudinal transcriptome studies have worked around this issue by inhibiting reproduction either through application of chemical inhibitors like Fluorodeoxyuridine (FUDR) (Ham et al., 2022; Li et al., 2019; Rangaraju et al., 2015), or by using strains with condition-specific induced sterility such as *rrf-3* and *fem* mutants (Barton et al., 1987; Garigan et al., 2002; Lund et al., 2002; Murphy et al., 2003). Although these are useful methods to ensure no progeny are mixed in with the adult population when extracting RNA, the practice does move the animals away from a natural condition, affecting the transcriptome (Anderson et al., 2016; Feldman et al., 2014). A recent study also showed that the amino acid biosynthesis of *E. coli* exposed to FUDR is altered, which can affect gene transcription of feeding nematodes (McIntyre et al., 2021). In the transcriptome generation for this study, we opted to not interfere with the process of reproduction. The number of cells contributing oocyte-related transcripts present in an actively reproducing hermaphrodite should be acknowledged, but is relatively small (Corsi et al., 2015; Schafer, 2005) and therefore unlikely to majorly affect the conclusions of our study. Other longitudinal transcriptome studies were performed under similar conditions (Golden et al., 2008; Schmeisser et al., 2013; Tarkhov et al., 2019).

In conclusion, we have shown the occurrence of a reproductive shift in the transcriptome of *C. briggsae*. Looking into this, we found that the genes affected by this shift are highly represented by several biological processes, reproduction most determinedly. Additionally, our findings reveal that genes involved in transcription and regulation of gene expression are downregulated post-reproductively. One possibility of such a change may be an overall reduction in gene regulation in older adults as is predicted by the run-on effect of a quasi-program (Gems and Kern, 2022). Presenting the first adult longitudinal transcriptome in *C. briggsae*, we believe our data and analysis will provide a valuable resource to the field, allowing for more in-depth study of this species and of *Caenorhabditis* nematodes at large. Additionally, the work sets the stage for future research into the conserved mechanisms governing reproductive biology and aging and their broader implications across different animal models.

## ACKNOWLEGEMENTS

We thank members of the Gupta lab for discussions throughout this project. We would like to acknowledge Avijit Mallick for assisting with sample preparation and being instrumental in initial planning, Nikita Jhaveri for assisting with initial RNA-seq analysis, Ron Ellis for suggesting altered reproduction experiments and providing the *Cbr-glp-4(v273)* strain, and Paul Sternberg for comments on the manuscript. This work was supported by the Natural Sciences and Engineering Research Council of Canada (NSERC) Discovery grant to BPG.

## SUPPLEMENTAL MATERIALS

### Supplemental Results

#### Gene ontology analysis

The matrisome is the matrix of proteins and connective material that sits in between cells (Teuscher et al. 2019), and undergoes dynamic changes during the life of animals (Candiello et al. 2010, Keeley & Sherwood, 2019). This includes processes involved with the extracellular matrix (ECM), basement membrane, cuticle and collagens (Supplemental file 3, Supplemental figure S1). The post-reproductive set were represented by genes belonging to col, bli, dpy and sqt classes, in such terms as ‘collagen and cuticulin-based cuticle development’, ‘collagen and cuticulin-based cuticle extracellular matrix’, and ‘cuticle development’.

Our analysis showed some interesting examples of basement membrane genes that undergo expression changes between reproductive and post-reproductive phases. For example, *Cbr-pat-3* and ECM hemicentin *Cbr-him-4* have constant expression levels during reproduction but are upregulated after. Together with *Cbr-ina-1* and *Cbr-vab-10*, these genes are involved in structural integrity of the body, serving to bind together basement membranes to withstand the physical stresses such as egg laying (Morrissey et al. 2014). However, *Cbr-ina-1* and *Cbr-vab-10* do not become overexpressed during the postreproductive day 6-9, suggesting that instead of egg-laying stress causing the expression patterns seen for *Cbr-him-4* and *Cbr-pat-3,* it is more likely due to other processes such as general tissue maintenance. The basement membrane, particularly laminins within it, in adult nematodes has a function to maintain proteostasis in muscle cells, and has been implicated in neuro-muscular integrity (Jensen et al. 2012).

Matrisome genes are also involved in lifespan regulation. Teuscher et al (2019) reported an enrichment for collagens and other ECM genes among those involved in longevity (79 of 426 genes). We found that 60 of the 79 genes have orthologs in *C. briggsae*, and 25 of these follow the down-invert pattern, which represents a significant proportion (hypergeometric test p<0.001).

Furthermore, we found enrichment for terms related to axonal guidance. Axon guidance factors (netrins, slits, agrin, fibrillin) are part of the basement membrane, in a manner conserved between nematodes and mammals (Teuscher et al. 2019). Although the process of axonal guidance is most known for its role in development (Yurchenco & Wadsworth 2004; Sherwood 2021), their role continues in adult life for neuronal repair and regeneration (Roet & Verhaagen, 2014). While in *C. elegans* there was no enrichment for related GO terms in the day 1-6 set, we observed enrichment among postreproductive upregulated genes. In *C. briggsae,* axonal guidance terms are down-invert for 1-6 and 6-9 with terms related to biological processes, i.e. ‘axon guidance’, and for cellular component terms including ‘axon’ and ‘neuron projection membrane’. Representative genes found include the type XVIII collagens *Cbr-cle-1* and *Cbr-col-99*, of which the *C. elegans* orthologs are known to be involved in axon guidance (Ackley et al. 2001; Taylor et al. 2018).

Many of the terms in the up-invert DNA damage repair category were related to DNA replication. The examples included ‘DNA synthesis involved in DNA repair’, with DNA polymerase eta and kappa (*Cbr-polh-1*, *Cbr-polk-1*), alongside ‘regulation of DNA replication’ with genes having no apparent role in DNA damage repair such as *Cbr-cyd-1*/cyclin D, DNA polymerase facilitators *Cbr-pcn-1* and *Cbr-cdt-1*, and the DRM complex component *Cbr-lin-35* involved in many developmental processes (González-Rangel & Navarro, 2017). The post-reproductive enrichment includes ‘DNA repair’ with DNA replication factors *Cbr-rfc-2* and *Cbr-rfc-3.* In *C. elegans, rfc-3* protects DNA from damage (Pothof et al. 2003). Lastly, ‘UV-damage excision repair’ is well-represented, with *Cbr-xpa-1, CBG23450 and Cbr-ercc-1* i.e. all genes annotated for the process. Similar to GO, the KEGG analysis also revealed enrichment for DNA damage repair pathway (namely, *parp-1, pcn-1, csb-1* and *mlh-1*). Indeed, after the reproductive phase, upregulated genes show only negative enrichment for KEGG pathways DNA damage repair and DNA replication, supporting the hypothesis that after day 3 DNA damage repair declines. Branching from DNA damage repair are the additional enriched KEGG terms Mismatch repair, Nucleotide excision repair and Non-homologous end joining (Supplemental file 2). What seems most likely is that DNA repair and synthesis factors are highly expressed early on as part of active reproduction, where DNA is constantly being produced in the developing oocytes and embryos present inside the body. DNA synthesis requires simultaneous DNA repair activity as the process has a significant rate of introducing errors during replication.

Additional supporting data for lipid metabolism annotation for the reproductive shift comes from the KEGG pathways ‘fatty acid metabolism’ and ‘fatty acid biosynthesis’. These are based on the genes *acs-5,16,18;* dehydrogenase *dhs-25* and oxydoreductase *Y48A6B.9.* The fatty acid metabolism is further enriched with *fat-5, acdh-1, acdh-18, ech-6*, and hydroxyacyl-CoA dehydrogenase *F54C8.1.* The *acs* genes are involved in fatty acid biosynthesis, and *ech* genes are essential for metabolizing fatty acids in beta oxidation to produce both acetyl CoA and energy in the form of ATP (Watson et al. 2016). Post-reproductively the same enrichment is still present, but only among upregulated genes, for ‘fatty acid metabolism’ with all the same genes except for *Y48A6B.9*.

Additional non-lipid catabolism terms came up for day 1-3, where enrichment was apparent for the term ‘small molecule catabolic process’, in *C. elegans* among up- and downregulated genes, but in *C. briggsae* only with downregulated genes. The term ‘carboxylic acid catabolic process’ featuring 6 hydrolases including *Cbr-ahcy-1, Cbr-fah-1* and *Cbr-kynu-1* are downregulated. The term ‘protein catabolic process’ is negatively enriched among downregulated genes. Many of these processes all come up the similarly when using either the original *C. briggsae* genes or *C. elegans* orthologs. However, species-specifically we also see ‘alcohol catabolic process’ enriched only for *C. elegans*, and several terms for *C. briggsae* including ‘peptidoglycan and aminoglycan catabolic processes’. Other affected terms included ‘protein catabolic process’ that was enriched among reproductively upregulated genes, ‘proteasomal protein catabolic process’ and ‘proteasome-mediated ubiquitin-dependent protein catabolic process’ were enriched among post-reproductively downregulated genes.

Many GO terms involved in gene transcription and the regulation thereof were enriched. The analysis of day 1-3 upregulated genes revealed enrichment for terms of negative as well as positive regulation of transcription: ‘negative regulation of transcription by RNA polymerase II’ ‘negative regulation of DNA-templated transcription’ and ‘positive regulation of DNA-templated transcription’. Post-reproductively, the downregulated genes registered with positive and negative regulation of transcription based on *C. briggsae* annotation. Among the relevant terms were ‘negative regulation of DNA-templated transcription’, and ‘positive regulation of DNA-templated transcription’. Genes exclusively involved in positive regulation of transcription included *Cbr-glp-1, Cbr-sea-1.2, Cbr-sea-1.4* and *Cbr-lag-1.* From studies in *C. elegans*, the orthologous *sea-1/sea-2* is known to be a transcription factor involved in sex determination during early development (Powell et al. 2005; Huang et al. 2011), and *glp-1* and *lag-1* are involved in Notch signaling. Among the genes exclusively involved in negative regulation are *Cbr-flh-1* and *Cbr-flh-3*, which are involved in repression of microRNA genes during development (Ow et al. 2008). It is likely that expression of these factors during reproduction, and downregulation thereafter is either signal from the reproducing mother, or the eggs in various stages of development inside the uterus of measured animals.

## SUPPLEMENTAL TABLES

**Supplemental Table 1.**
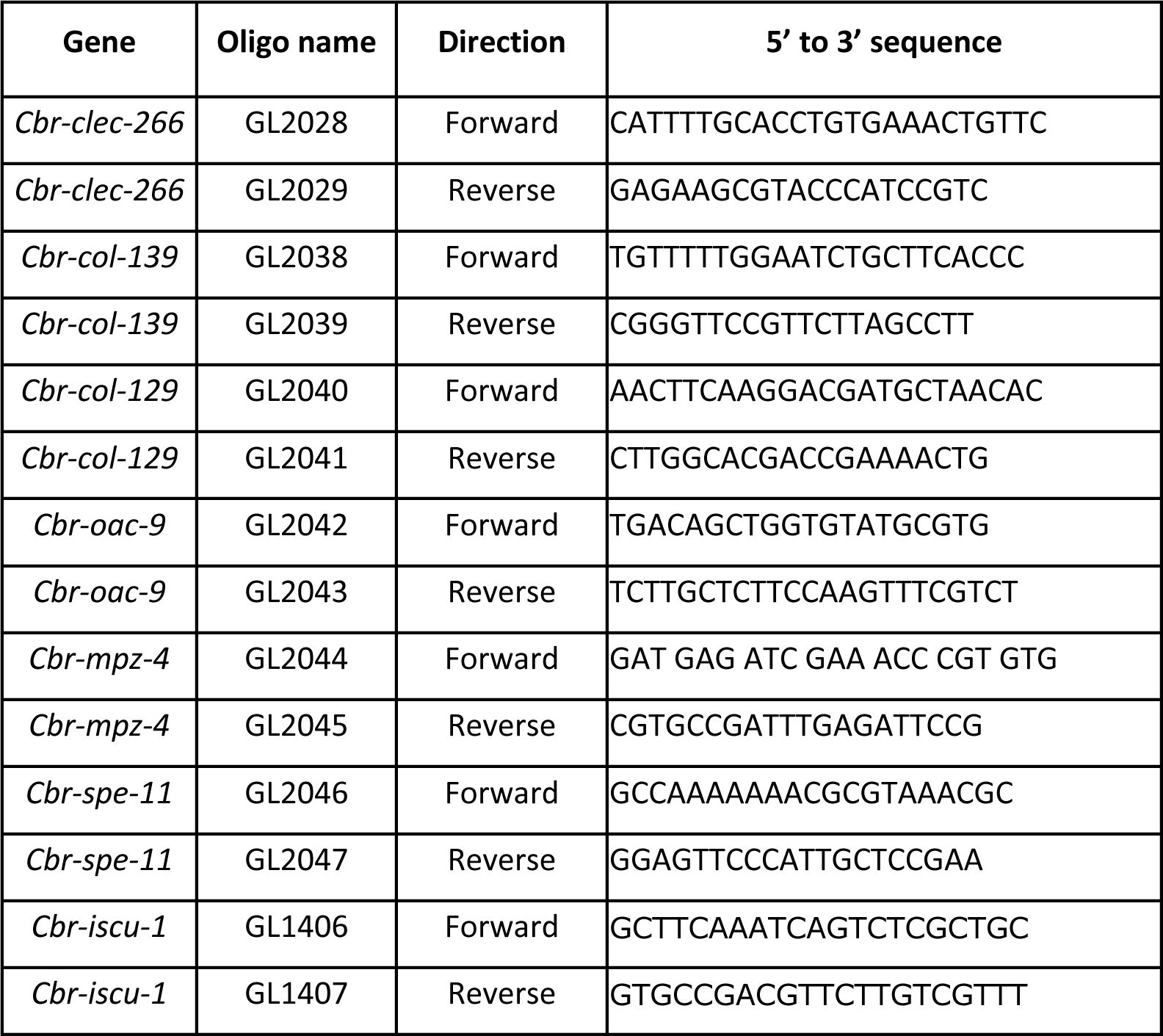
Primers used in qPCR experiments.

**Supplemental Table 2.**
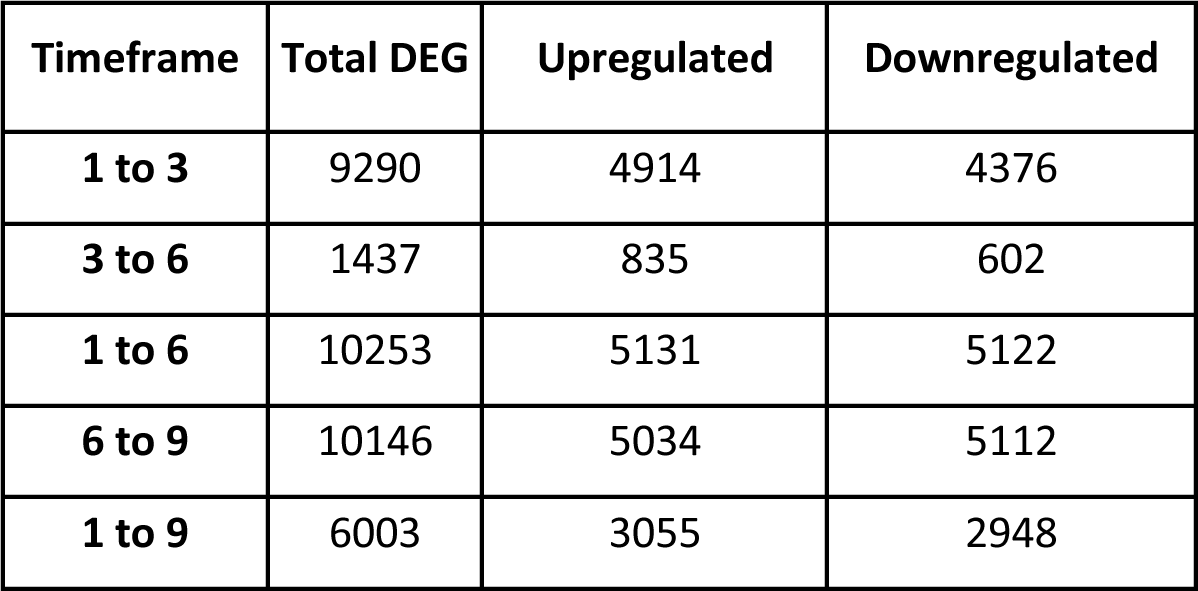
Age-based differentially expressed *C. briggsae* genes without fold change cutoff. Numbers of DE genes (p<0.01) obtained from pairwise comparison of RNA-sequencing on differently aged synchronized *C. briggsae* populations. The table contains numbers of all genes, without applying a minimum 2-fold expression change cutoff.

| Timeframe | Total DEG | Upregulated | Downregulated |
| --- | --- | --- | --- |
| **1 to 3** | 9290 | 4914 | 4376 |
| **3 to 6** | 1437 | 835 | 602 |
| **1 to 6** | 10253 | 5131 | 5122 |
| **6 to 9** | 10146 | 5034 | 5112 |
| **1 to 9** | 6003 | 3055 | 2948 |

**Supplemental Table 3.** Age-based differentially expressed *C. elegans* genes. DE genes (p<0.01) obtained from pairwise comparison of RNA-sequencing on differently aged synchronized *C. elegans* populations based on transcriptome data from Schmeisser et al. (2013). The table contains numbers of genes after applying a minimum 2-fold expression change cutoff (log2fold expression +/−1).

| Timeframe | Total DEG | Upregulated | Downregulated |
| --- | --- | --- | --- |
| 1 to 5 | 3535 | 935 | 2600 |
| 5 to 10 | 1137 | 476 | 661 |
| 1 to 10 | 3983 | 1338 | 2645 |

**Supplemental Table 4.** Gene expression in reproduction-impaired *Cbr-glp-4(v273)* and reproduction-extended mated AF16. Expression and significance values for genes used in *Cbr-glp-4(v273)* and mated AF16 experiments.

| Gene name | Expression change between timepoints |  |  |  |  | adjusted p values |  |  |
| --- | --- | --- | --- | --- | --- | --- | --- | --- |
|  | day 1-3 | day 3-6 | day 1-6 | day 6-9 | day 1-9 | adj. p day 1-3 | adj. p day 3-6 | adj. p day 6-9 |
| *Cbr-clec-266* | 3.12 | -0.55 | 2.03 | 1.38 | 3.32 | 1.74E-03 | 1.50E-02 | 3.47E-13 |
| *Cbr-col-139* | -5.72 | -0.42 | -8.59 | 6.09 | -2.34 | 2.9E-168 | 6.03E-52 | 4.5E-103 |
| *Cbr-col-129* | -5.64 | -0.38 | -7.95 | 5.41 | -2.40 | 0 | 7.9E-120 | 4.51E-116 |
| *Cbr-oac-9* | -5.24 | -0.41 | -6.61 | 4.31 | -2.14 | 1.19E-73 | 4.29E-20 | 5.67E-48 |
| *Cbr-mpz-4* | -5.21 | -0.02 | -5.13 | 3.41 | -1.01 | 6.23E-09 | 3.35E-03 | 2.39E-06 |
| *Cbr-spe-11* | -4.90 | -0.61 | -7.62 | 5.01 | -2.49 | 5.65E-152 | 6.52E-46 | 3.64E-127 |

## SUPPLEMENTARY FIGURES

**Supplemental Figure S1:**
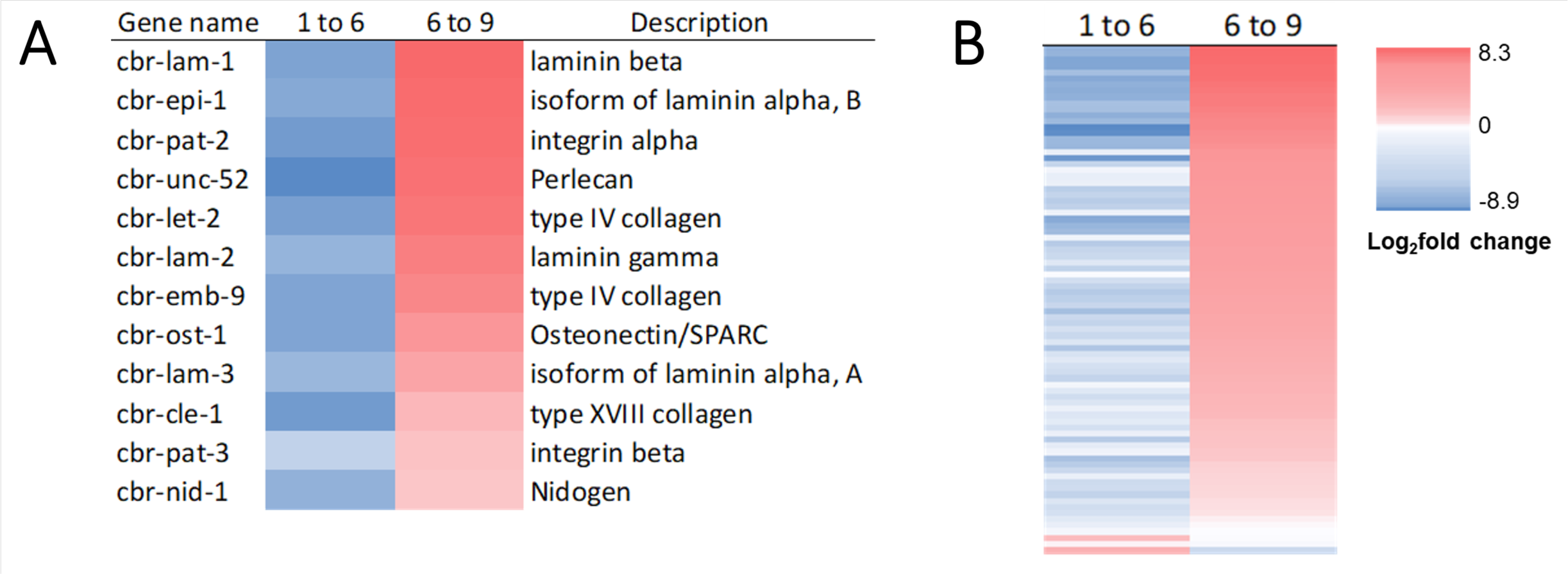
Heatmaps of differentially expressed matrisome genes. **A.** Basement membrane genes, day 1 to 6 and 6 to 9. **B.** Cuticular collagens, day 1 to 6 and 6 to 9.

**Supplemental Figure S2:**
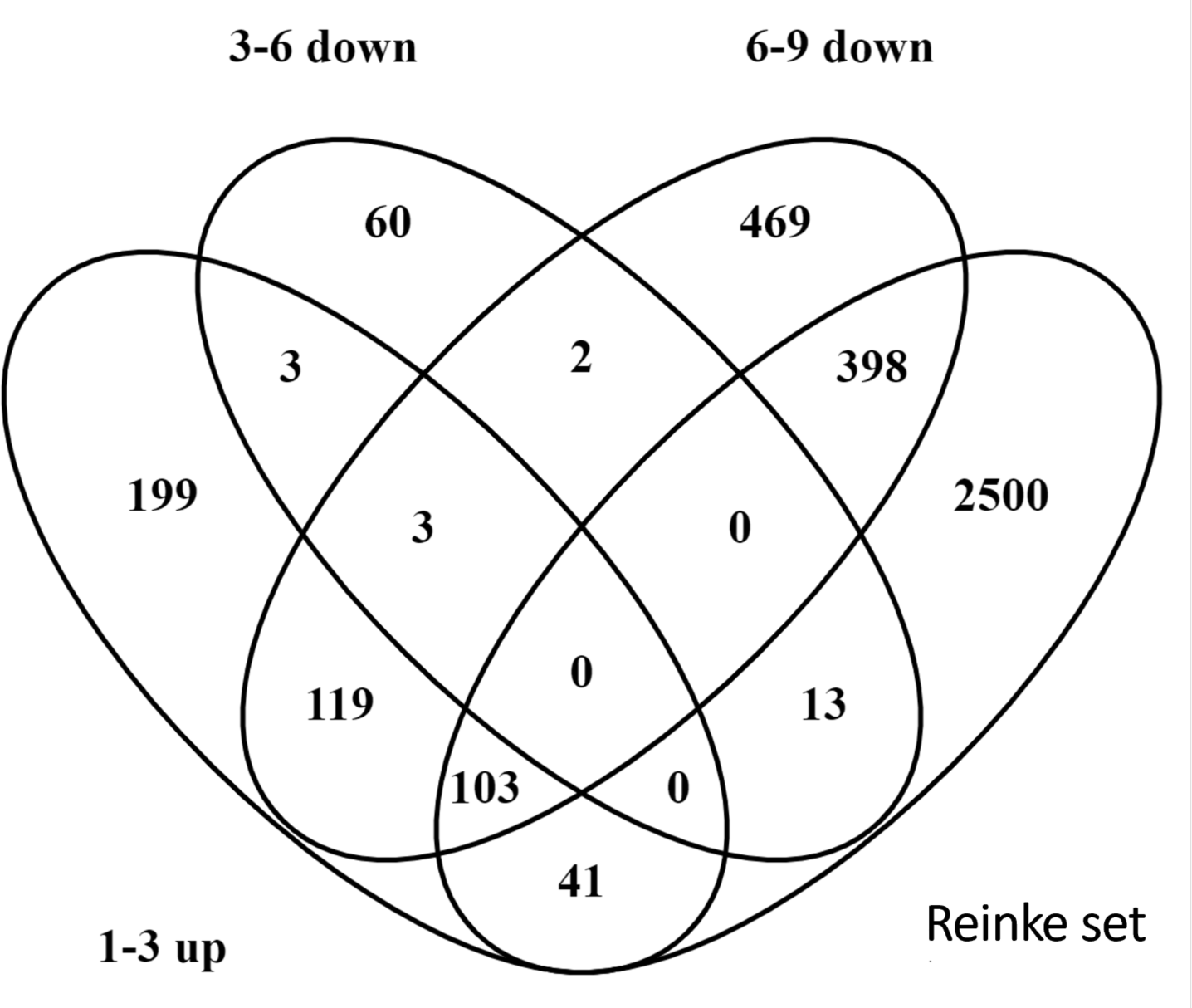
Overlap *C. briggsae* DEGs and canonical germline genes. Venn diagram of overlap between *C. elegans* 1- to-1 orthologs of *C. briggsae* DEG’s at 3 timeframes, and germline gene set by Reinke et al. (2004).

**Supplemental Figure S3:**
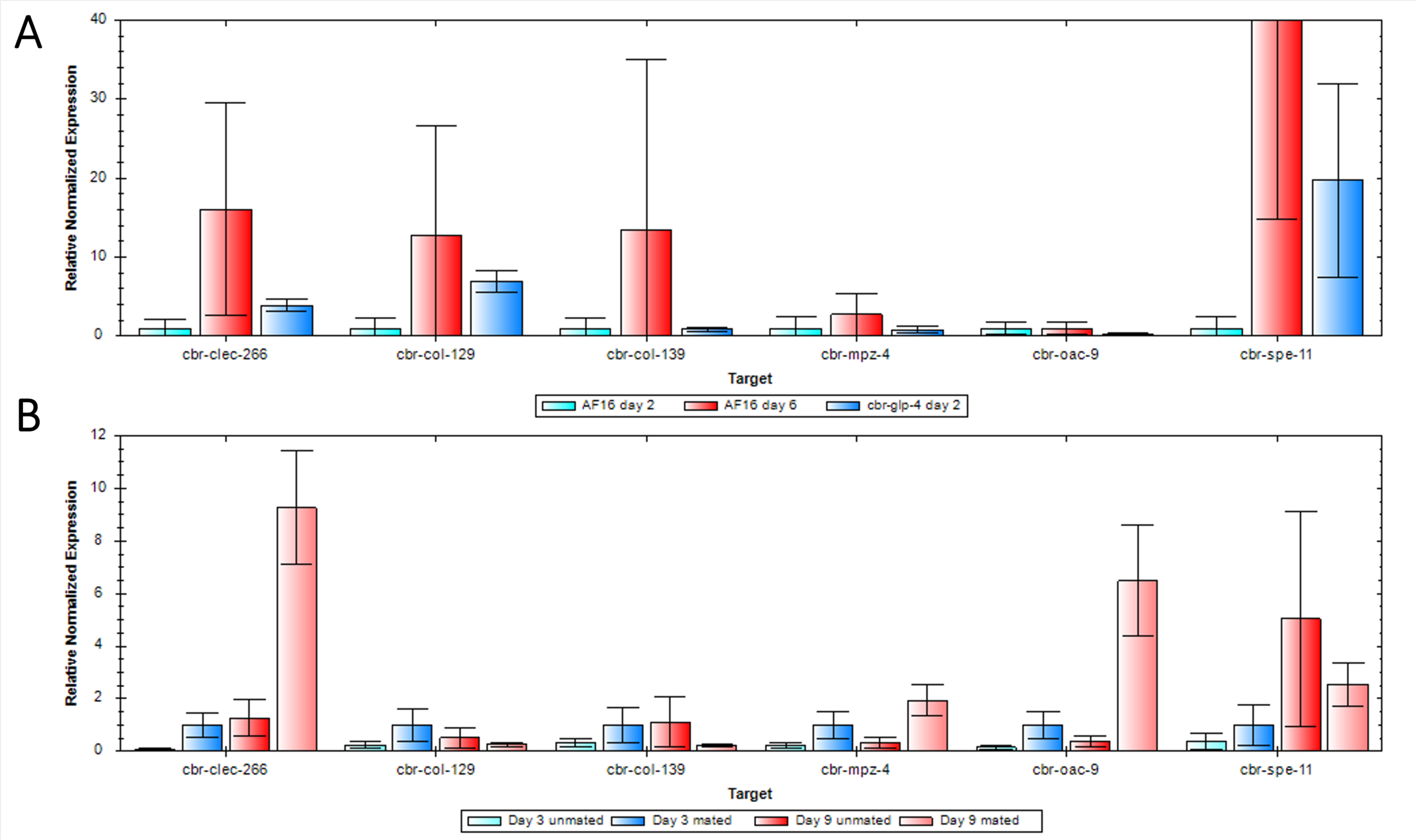
Gene expression in reproduction impaired Cbr-glp-4(v273) and reproduction extended mated AF16. **A.** Experiments performed at 26C in *C. briggsae* AF16 and *Cbr-glp-4(v473). Cbr-clec-266* is involved in immunity, *Cbr-col-129 & Cbr-col-139* in cuticle formation and *Cbr-mpz-4, Cbr-oac-9* and *Cbr-spe-11* in germline processes. *Cbr-spe-11* AF16 day 6 is 76.3 +/− SEM 61.6. Values are normalized to AF16 day 2. **B.** Animals reared at 20℃, aged day 3 and 9 of adulthood. Mating occurred from day 7 to 8 of adulthood. Values are normalized to AF16 day 3 mated samples.

## Notes

### Competing Interest Statement

The authors have declared no competing interest.

